# High-resolution phenotypic landscape of the RNA Polymerase II trigger loop

**DOI:** 10.1101/068726

**Authors:** Chenxi Qiu, Olivia C. Erinne, Jui Dave, Ping Cui, Huiyan Jin, Nandhini Muthukrishnan, Leung K. Tang, Sabareesh Ganesh Babu, Kenny C. Lam, Paul J. Vandeventer, Ralf Strohner, Jan Van den Brulle, Sing-Hoi Sze, Craig D. Kaplan

**Author notes:** Correspondence should be addressed to C.D.K.

## Abstract

The active site of multicellular RNA polymerases have a “trigger loop” (TL) that multitasks in substrate selection, catalysis, and translocation. To dissect the *Saccharomyces cerevisiae* RNA polymerase II TL at individual-residue resolution, we quantitatively phenotyped nearly all TL single variants *en masse*. Three major mutant classes, revealed by phenotypes linked to transcription defects or various stresses, have distinct distributions among TL residues. We find that mutations disrupting an intra-TL hydrophobic pocket, proposed to provide a mechanism for substrate-triggered TL folding through destabilization of a catalytically inactive TL state, confer phenotypes consistent with pocket disruption and increased catalysis. Furthermore, allele-specific genetic interactions among TL and TL-proximal domain residues support the contribution of the funnel and bridge helices (BH) to TL dynamics. Our structural genetics approach incorporates structural and phenotypic data for high-resolution dissection of transcription mechanisms and their evolution, and is readily applicable to other essential yeast proteins.

---

RNA polymerase II (Pol II) synthesizes all eukaryotic mRNA. Structural studies of *Saccharomyces cerevisiae* (*Sce*) Pol II have illuminated mechanisms of transcription^1-6^, especially RNA synthesis. RNA synthesis occurs through iterative nucleotide addition cycles (NACs): selection of correct substrate nucleoside triphosphate (NTP), catalysis of phosphodiester bond formation, and enzyme translocation to the next template position. These critical steps in NAC appear to be coordinated by a critical, conserved domain within the Pol II active site: the trigger loop (TL).

TL functions are underpinned by its mobile and flexible nature (Fig. 1a). The primary function of TL is kinetic selection of correct NTP substrates while balancing transcription speed and fidelity, and this function is highly conserved based on studies of RNAPs from *E. coli* (*Eco*)^7,8^, *T. aquaticus* (*Taq*)^9^, the archaeon *P. furiosus* (Pfu)^10^, and eukaryotic Pol II from *Sce*^11,12^ and human^13^. In a simplified two-step model, correct NTP binding appears to facilitate TL movement such that a bound, matched NTP shifts the TL from the “open” state to the “closed” state^4,14-17^, allowing capture of the matched NTP in the Pol II active site and promotion of phosphodiester bond formation^4,16,18^. The subsequent release of the byproduct, pyrophosphate, allows a conformational shift of the TL from the “closed” state back to the “open” state^14,19,20^. TL opening has been proposed to be critical for enzyme translocation relative to template, an essential step for the next nucleotide addition cycle^8,12,14,21-24^. Furthermore, additional TL states have been implicated in transcriptional pausing from studies in *E.coli*^16,21,25^, backtracking from structural observations^26,27^, and, although controversial, intrinsic cleavage^7,28-31^. Thus, distinct TL conformations or interactions are linked to different functions in transcription, with delicate control of TL dynamics promoting proper transcription elongation while possibly incorporating signals from the rest of Pol II or Pol II bound factors^16,32-35^.

**Figure 1:**
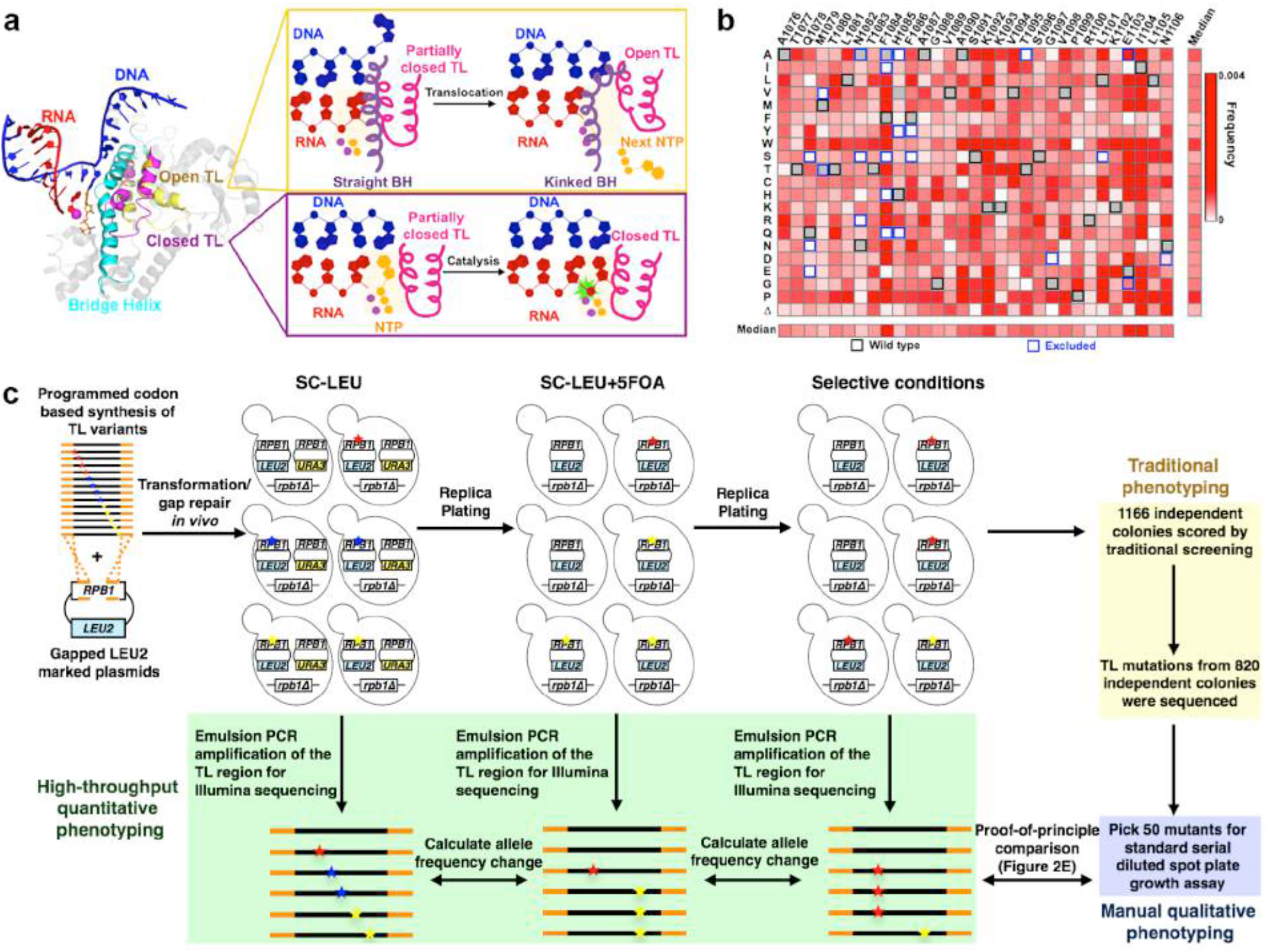
Establishment of a high-throughput platform for phenotyping comprehensive TL single variant library. Figure 1a: Multiple TL functions are underpinned by its mobile nature. Structures of open (PDB:5C4J) and closed TLs (PDB:2E2H) are shown in the context of surrounding domains. DNA (blue), RNA (red), Bridge Helix (cyan), Closed TL (magenta) and Open TL (yellow) are shown in cartoon rendering. The open TL has been proposed to allow Pol II translocation while the closed TL has been shown to facilitate catalysis (right panel). Figure 1b: Mutational coverage of the TL variant library is shown as heatmap illustrating the allele frequencies of single substitution variants (WT amino acids and positions labeled on x axis; amino acid substitutions on y axis). The WT amino acids for each position are noted with black boxes, and mutants excluded from library synthesis are noted using blue boxes. Figure 1c: Schematic representation of experimental approach. Stars of different colors represent distinct substitutions. The TL variant library PCR amplicon (encoding Rpb1 amino acids 1076-1106) flanked by *RPB1 TL* flanking sequence (orange) was co-transformed with a linearized *LEU2 CEN* plasmid containing an *rpb1* gene with the TL deleted, allowing construction of full-length *RPB1* (with TL variants) by *in vivo* homologous recombination. Heterozygous Leu^+^ transformants were replica-plated onto SC-LEU+5FOA to select against the WT *RPB1* (*URA3 CEN*) plasmid and to create TL variant pools. TL variant pools were subsequently replica-plated to different selective conditions for either traditional individual colony screening or high-throughput phenotyping using deep sequencing. For the latter, replica-plated colonies were pooled for genomic DNA extraction, and the TL region was amplified by emulsion PCR to prepare templates for deep sequencing.

Genetic and biochemical studies have revealed TL functions in the NAC. First, the nucleotide interacting region (NIR, Rpb1 1078-1085) discriminates matched rNTPs from 2’-dNTPs and non-complementary rNTPs^11,12^. NIR substitutions widely conferred lethality in residues observed to interact with rNTPs. Where viable, substitutions reduced catalytic activity *in vitro* and were termed as partially loss-of-function (LOF)^7-11,36^. Second, a TL C-terminal mutant E1103G, conferred increased catalytic activity *in vitro*, which we termed gain-of-function (GOF)^11,12,37^. Fast kinetics experiments revealed that E1103G may bias TL dynamics towards the catalytically active“closed” state^12^, consistent with infidelity and compromised translocation in addition to increased catalysis^11,12,22,38,39^. Furthermore, we previously described a set of Pol II TL mutants with broad and distinct alterations to transcription *in vivo*, thus conferring allele-specific phenotypes (Table 1) that correlate with decreased or increased activity^36,40^ *in vitro*. Various genetic interactions (suppression, exacerbation, and epistasis) have also been observed, suggesting a complex functional network within the Pol II TL^36^. Finally, context dependence for TL function was observed, wherein analogous mutations in a conserved TL residue showed opposite effects in *Sce* Pol I and Pol II, suggesting different rate limiting steps for the two enzymes^41^. Together, the intricate intra-TL functional network and the context dependence of TL properties suggest importance of extensive residue-residue interactions within or outside the TL.

**Table 1.**
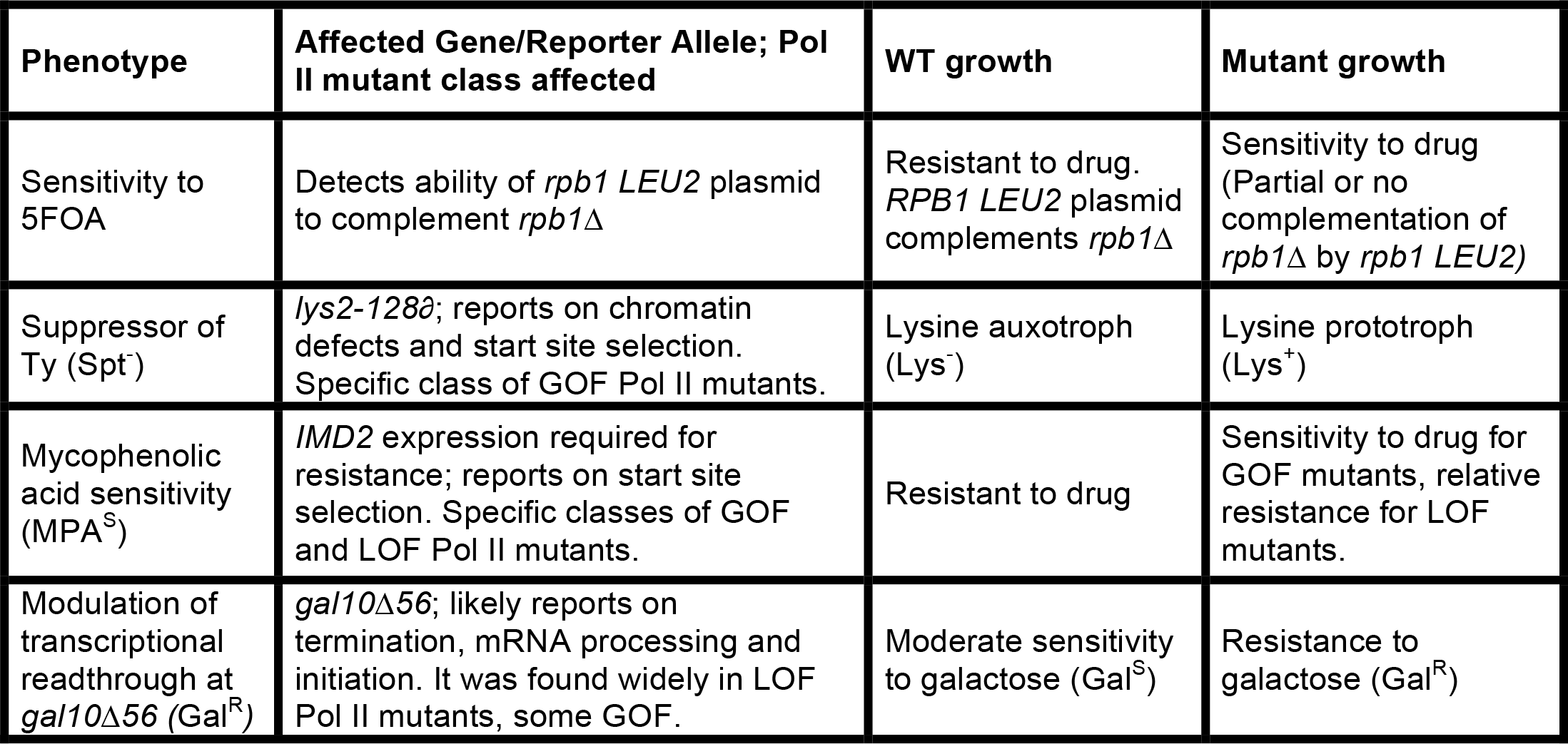
Plate phenotypes employed for the screening Pol II alleles in vivo.

The possible multifunctional nature of each TL residue complicates interpretations of functions based on a limited number of mutants, since the phenotype of any given mutant could result from removal of the wild type side-chain or additional functions of the substituted one. Furthermore, different substitutions may have distinct effects on particular TL conformations^36,42^. In the TL, different substitutions in the same residue can confer distinct phenotypes, so limiting mutational analyses to a single substitution at a particular position may mislead about residue function^12,36^. Deep mutational scanning is an emerging technique for studying large sets of mutants by assessing the enrichment or depletion of variants after a strict selection process^43^. Different selection approaches have been designed such that a specific protein property (sensitivity to substitutions^44^, thermo stability^45^, protein stability^46^, *etc*) can be studied. Notably, our established genetic phenotypes (Table 1) were well correlated with altered transcription elongation rates *in vitro* and multiple specific transcription defects *in vivo*^36,40^, thus providing a powerful phenotypic framework for studying TL function. In this work, we have defined the phenotypic landscape of the conserved, essential *S. cerevisiae* Pol II TL. We have found three major, distinct classes of transcriptionally defective TL mutants associated with differential stress response profiles, and determined the functional contributions of TL residues. We have interrogated the mechanisms by which proximal Pol II domains communicate with the TL, while identifying examples of epistasis between residues, which are the likely drivers of incompatibility of RNAP evolutionary variants when placed in the Pol II context.

## Results

### Strategy for studying *in vivo* effects of TL variants library

A comprehensively mutagenized TL variants library (Rpb1 1076-1106), with well-characterized TL mutants^11,36^ excluded for further controls, was synthesized using the Slonomics technology^47,48^ and validated by deep sequencing (Fig. 1b). Synthesis conditions were such that single substitution mutants would predominate. Our TL mutant library showed even distribution of substitutions across all positions and substitution types (Supplementary Fig. 1a-c), with generally very low frequencies for excluded mutants (Fig. 1b). We first sought evidence that measured allele frequencies reflected the real distribution of mutants, because PCR fidelity for similar amplicons is often compromised by template switching^49,50^. We spiked in five excluded single variants (H1085Y/Q, F1086S, G1097D, E1103G) as a control, so double mutants from the spike-in variants should not be present in our library but presumably the result of template switching. We prepared TL amplicons from a subset of conditions using both standard PCR and emulsion PCR (emPCR), which can suppress template switching^49,50^. First, double mutants from spike in controls were found at a significantly lower frequency than relevant single variants, and second, emPCR further suppressed the template switching frequency for all possible double mutants derived from spike-in single variants (Fig. 2a, left) 2.5 fold on average (Fig. 2a, right). We conclude that template switching is likely not extensive in our reactions but further reduction by emPCR led us to employ emPCR for our studies.

**Figure 2:**
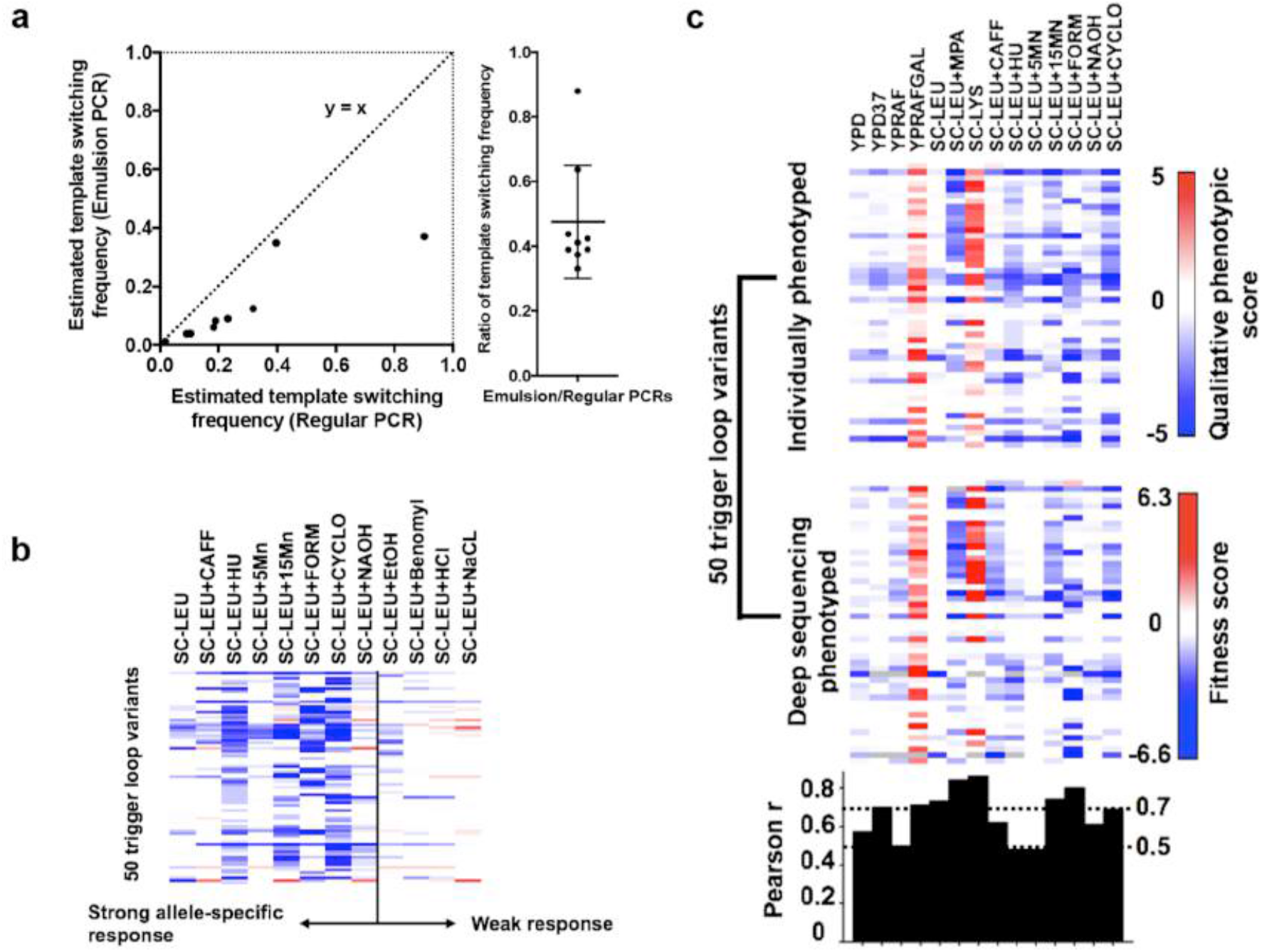
Quality controls for the high-throughput phenotyping approach. Figure 2a: Comparison of estimated template switching frequency in regular and emulsion PCR conditions. Template switching was estimated by the ratio (Freq^Double^) / (Freq^Single1^ × Freq^Single2^) for all the possible double mutants combined from five spiked-in single mutants. Figure 2b: Additional growth conditions were employed to increase resolution for distinguishing similar TL alleles. Growth scores for 50 individually isolated TL mutants (y axis) under 12 growth conditions (x axis), as determined by standard serial dilution plate phenotyping, are shown as a heatmap (Supplementary figure 2). Positive values shown in red indicate better growth than WT and negative values in blue indicate worse growth than WT. Figure 2c: High-throughput quantitative phenotyping results are consistent with individual phenotyping of variants. Top heatmap shows qualitative growth scores (as in Fig. 2b) of 50 individually phenotyped TL variants on y axis (supplementary figure 2) with selective conditions on the x axis. Deep sequencing results for the same mutants using median of fitness defects from three independent high-throughput screens are shown in the middle panel. Pearson r calculated to show the correlation between each condition from the two datasets were shown in the bottom panel.

We have developed an experimental pipeline to examine mutations in an essential gene using a plasmid shuffling strategy, and have applied it to phenotyping the TL variant library (Fig. 1c). To validate our pipeline and to isolate novel TL alleles for further controls, we performed a traditional genetic screening for mutants with transcriptional defects (Table 1) and isolated 1166 candidate mutants (Supplementary Table 1), which included 154 singly-substituted and 386 multiply-substituted variants. To further distinguish mutants, we examined 50 single variants under various stress conditions (Fig. 2b, Supplementary Fig. 2a), and observed that media containing caffeine, hydroxyurea, MnCl_2_, formamide, cycloheximide, or NaOH induced allele-specific sensitivity or resistance, while media containing ethanol, benomyl, HCl or NaCl showed fewer allele-specific effects (Fig. 2b, Supplementary Fig. 2a). Therefore, in our high-throughput approach, we phenotyped TL variants library under our established conditions (Spt^-^, MPA^S^, Gal^R^) and stress conditions exhibiting allele-specific discrimination ability. Phenotypic score was estimated from the change of allele frequency normalized to WT, as is standard for previous mutational scanning studies^43-46^. Quantitative phenotypic scores of the 50 mutants from the high-throughput phenotyping were consistent with semi-quantitative growth scores derived from standard phenotyping (Fig. 2c, supplementary Fig. 2), validating our approach.

### The Pol II TL fitness landscape

The TL is highly conserved, especially in the NIR, the loop tip residue (Rpb1 G1088) and several TL C-terminal residues (Fig. 3a). Highly conserved residues are predicted to be critical for protein function, thus substitutions are expected to confer fitness defects and be selected against. We first sought to evaluate general fitness defects of observed TL singly-substituted variants (termed “fitness landscape”), both in the presence of WT *RPB1* (Fig. 3b) and upon removal of WT *RPB1* (Fig. 3c). Notably, TL NIR and loop tip substitutions conferred large fitness defects in general, while most perturbations in the similarly conserved C-terminal residues did not confer severe growth defects (Fig. 3b, 3c). This observation highlights that conservation does not necessarily reflect sensitivity to perturbations, and TL fitness landscape can further distinguish extremely highly conserved TL residues, as discussed below:

**Figure 3:**
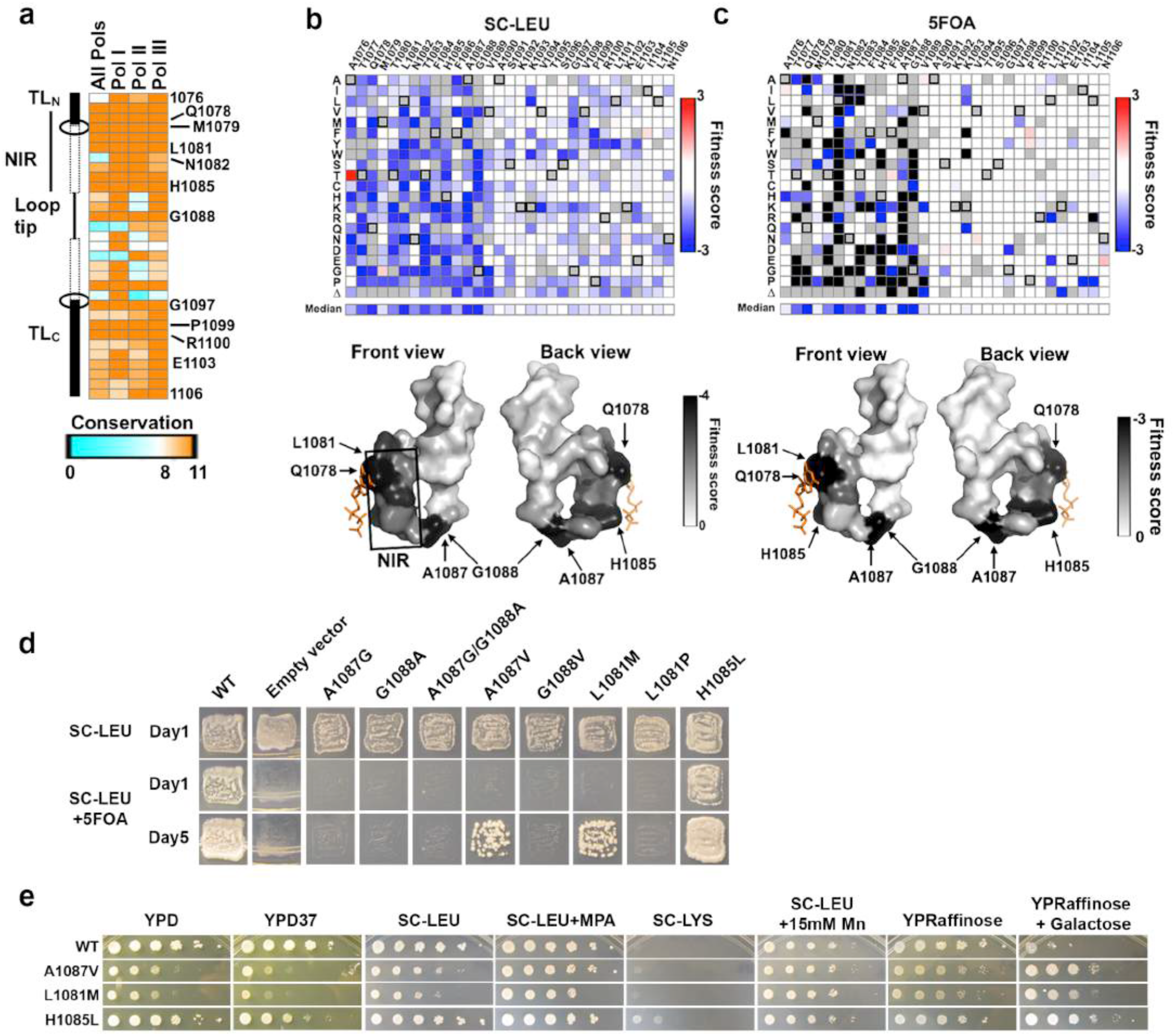
The TL fitness landscape distinguishes highly conserved TL residues and reveals high mutational sensitivity in the nucleotide interacting region (NIR) and the Alanine-Glycine linker. Figure 3a: Conservation heatmap of TL residues in eukaryotic RNA polymerases. The conservation scores were extracted from a multiple sequence alignment, including 182 Pol II, 59 Pol I, and 111 Pol III sequences. Figure 3b: Fitness defects of TL variants in the heterozygous state are shown as a heatmap. Unavailable data points are in grey squares. WT residues at indicated positions are in black box. Surface representation (bottom panel) of the TL structure (PDB:2E2H) is shaded by the median fitness value for all available variants at each position, in a gradient of white (rare defects) to black (common defects). The position of matched GTP substrate is shown in orange stick representation. Figure 3c: General fitness defects of TL variants upon removal of WT *RPB1*. Fitness defects predicted to result in lethality shown in black. Surface representation (bottom panel) of the TL structure is shaded by the median fitness value of all available variants at each position, in a gradient of white (rare defects) to black (defects common). Figure 3d: Complementation abilities of variants in the difficult-to-substitute TL positions (L1081, A1087, G1088) or unexpected TL variants (H1085L) assayed by plasmid shuffling of individual strains. Ability to grow on SC-LEU+5FOA indicates complementation of essential functions of *RPB1*. SC-LEU medium is the control state where WT *RPB1* is present. Figure 3e: Transcription-linked phenotypes of the viable mutants from the difficult-to-substituted residues L1081M, A1087V or unexpected TL variant H1085L.

First, substitutions in the NIR (Rpb1 1077-1085) generally conferred both fitness defects (Fig. 3c) and apparent dominance (Fig. 3b). Observed fitness defects were consistent with previous observations that several NIR mutants render Pol II slow in elongation *in vitro* and cause fitness defect *in vivo*^11,36^. Slowly elongating Pol II may engender clashes with WT Pol II on genes *in vivo*, consistent with the observed dominance in most NIR variants, and consistent with TL variants being assembled into Pol II complexes to interfere with WT Pol II function. Second, substitutions within the alanine-glycine linker (Rpb1 1087-1088) almost universally conferred lethality or severe growth defects. A Pol II structure with closed TL^4^ reveals that A1087 and G1088 are in a tight pocket between the funnel and bridge helices, presumably necessitating small side-chain residues (Supplementary Fig. 3a). To determine the extent of spatial constraint, we individually assessed the fitness of AG swapping variants and small hydrophobic valine substitutions (Fig. 3d). Notably, all the swapping variants (A1087G, G1088A and A1087G/G1088A) were lethal (Fig. 3d). While G1088V is lethal, A1087V is severely sick but viable (Fig. 3d), suggesting extremely high spatial constraint but differential tolerability for the two residues. This pocket/TL interaction is only observed in the closed TL^4^ but not in any of the open states^51^, suggesting function in stabilizing the active, closed TL conformation for promoting catalysis. Consistent with disruption of the pocket/TL interaction and the closed TL state, we observed LOF phenotypes for A1087V (Gal^R^, slight MPA^R^) (Fig. 3e). Finally, substitutions in the conserved C-terminal helix, though not strongly defective in general fitness, might have transcription defects and were further characterized (discussed below).

### Novel TL NIR mutants allow mechanistic insights

The TL fitness landscape identified residues highly sensitive to perturbations, while also revealing variants in NIR residues previously known to be difficult to viably substitute. We highlight L1081 and H1085 as two examples. L1081 directly interacts with the base of matched NTPs^4^, and the equivalent residues in *Eco* and *Taq* RNAPs are important for substrate selection or catalysis^7,9^. L1081 is the most sensitive to perturbations among the hyper-conserved NIR. All previously tested L1081 variants were lethal^36^, though viable substitutions were identified for all other NIR residues. Furthermore, GOF allele E1103G can generally suppress lethal substitutions for most NIR residues, but could not for tested L1081 substitutions. In our TL fitness landscape, almost all L1081 variants were indeed predicted to be lethal based on our fitness threshold (Fig. 3c). L1081M conferred a severe growth defect, but was predicted to be just above the viable threshold (Fig. 3c). To validate this prediction, we constructed L1081M for direct analysis, and found that L1081M was indeed viable yet severely sick (Fig. 3d). Furthermore, L1081M conferred Gal^R^ and slight MPA^R^ phenotypes, consistent with other LOF mutants (Fig. 3e). Eukaryotic multi-subunit RNA Polymerases share a stringent evolutionary requirement for L at this TL position, while bacterial and archaeal lineages show both M and L variants. Consistent with evolutionary tolerance of variation within bacterial and archaeal lineages, the *Taq* RNAP M1238L variant shows near WT activity for substrate selection and catalysis *in vitro*^9^. These results highlight epistasis within *Sce* Pol II, and likely eukaryotic RNAP lineages, imposing a stringent requirement for Leucine.

H1085 interacts with the β-phosphate of the matched NTP^4^, and has been implicated in substrate selection, catalysis, intrinsic cleavage and PPi release^28,52^. We previously constructed several H1085 variants (A/N/D/F were lethal, K/R/W/Y caused severe growth defects, Q caused slight growth defect^11,36,40^), suggesting that some polar or positively charged residues, but not a hydrophobic phenylalanine or alanine, could partially complement loss of the histidine^36^. Here, we found that H1085L was viable and healthy in the fitness landscape (Fig. 3c), and validated it with individual mutant analysis (Fig. 3d). While H1085L conferred slight MPA^R^ and Gal^R^ phenotypes, consistent with other LOF mutants (Fig. 3e), it also conferred a slight Spt-defect, suggesting distinct defects from most other NIR mutants and all known LOF mutants. This observation alters our understanding of the likely bounds of active site chemistry (see discussion).

### There are at least three distinguishable TL mutant classes

The overall TL fitness landscape revealed the essentiality of almost all single substitution TL variants in standard growth medium, but could not indicate the nature of transcriptional defects, as we had previously found that both LOF and GOF alleles conferred growth defects. Therefore, we sought to determine the phenotypic outcome of the TL variants for the transcription-related Gal^R^, MPA^S^ and Spt-phenotypes as well as for a variety of stress conditions (termed “phenotypic landscape”).

Hierarchical clustering of the phenotypic landscape for 412 TL variants passing fitness filters revealed three mutant classes with distinct features (Fig. 4a, 4b). Class 1 mutants generally conferred strong Gal^R^ phenotype yet were Spt^+^, and in some cases were also slightly MPA^R^ relative to WT, consistent with previously characterized LOF mutants. We also identified high formamide sensitivity as a new signature phenotype for Class 1 mutants. Class 2 mutants showed generally weaker Gal^R^, slight formamide resistance, but do not confer strong phenotypes otherwise, representing a novel TL mutant class yet to be biochemically characterized. Class 3 mutants generally conferred Gal^R^, Spt^-^ and MPA^S^ phenotypes, consistent with previously characterized GOF mutants. Mn^2+^ hypersensitivity (Mn^S^) was correlated broadly with Spt^-^ and MPA^S^ phenotypes, suggesting a relationship among these phenotypes, and consistent with previous *in vitro* biochemical and *in vivo* phenotypic data for a subset of known GOF mutants^53,54^. Notably, our spike-in LOF (F1086S, H1085Q and H1085Y) and GOF mutants (E1103G and G1097D) co-clustered with Class 1 and Class 3 mutants, respectively.

**Figure 4:**
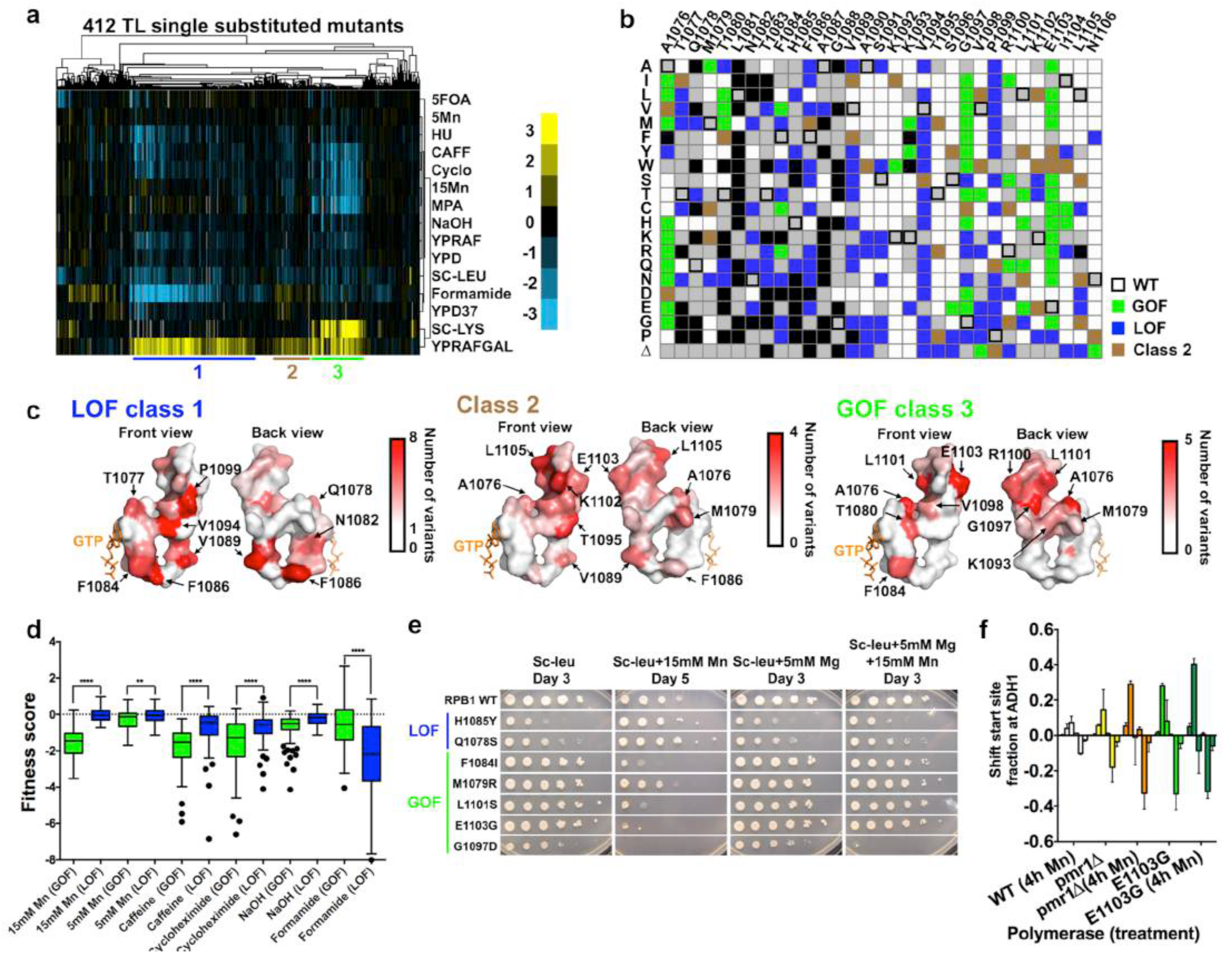
Three distinct TL mutant classes, revealed from TL phenotypic landscape, have specific distribution on the TL structure and distinct stress response profiles. Figure 4a: Hierarchical clustering of 412 single TL variants’ (y axis) phenotypes (calculated as in Figure 1c) under 14 different conditions (x axis) reveals distinct mutant classes. Positive (yellow) and negative (blue) fitness scores are shown as a heatmap. Mutant classes (clusters) are annotated by colored lines beneath the heatmap. Figure 4b: Distribution of three major mutant classes is shown in a single substitution variant heatmap. Class 1 (GOF) mutants are shown in green; Class 2 mutants are shown in brown and Class 3 (LOF) mutants are shown in blue. Figure 4c: Distribution of different mutant classes on the TL structure. TL is shown in surface and colored in the gradient from white to red by the number of clustered mutants at each position. Figure 4d: Differential stress responses in GOF and LOF mutants. GOF mutants are more sensitive to Mn, caffeine and cycloheximide, whereas LOF mutants are more sensitive to formamide. ^**^p<0.01, ^****^p<0.0001 (Two-tailed unpaired t-test). Figure 4e: Differential Mn^2+^ sensitivity and its suppression by Mg^2+^ for selected TL variants representative of mutant classes. Figure 4f: Mn^2+^ effects on different mutants’ transcription start sites (TSSs) distribution at ADH1, determined by primer extension analysis. TSSs at ADH1 are distributed in a range of positions and were divided into six bins for quantitation: from upstream (left) to downstream (right). Change of TSSs (normalized to untreated WT) is calculated by the fractions. Average and standard deviation of three experimental replicates are shown with plotted by bar graph with error bars.

### Functional contribution of TL residues in different states and substrate-induced TL closing mechanism

The distributions within different mutant classes predict distinct functional contributions of TL residues to TL dynamics. Perturbations predicted to bias the TL towards the active, closed TL state have been shown to result in GOF, whereas destabilization of the closed TL state generally leads to LOF^8,11,12,16,36^. Therefore, distributions of Class 1 (LOF) and Class 3 (GOF) mutants predict alterations to TL dynamics, as follows:

Class 1 (LOF) mutants included most variants from F1086, V1089, V1094 and P1099 (Fig. 4c, left), suggesting functions of these residues in stabilizing the closed TL. F1086 and V1089 are both proximal to multiple funnel helix residues when TL is closed^4,17^, while F1086 was proposed to orient H1085 for correct substrate interaction^17^. Therefore, alteration of these interactions may disrupt the closed TL state and result in LOF. Alternatively, recent Pol II structures revealed potential function of F1086-V1089 interaction in TL closing dynamics (Supplementary Fig. 3b). V1089 forms a backbone-backbone hydrogen bond with F1086 when TL is open, while its side chain flips towards the F1086 to form a hydrophobic interaction when TL is partially closed, suggesting that this side-chain interaction may be important for particular TL states (Supplementary Fig. 3b), though it was not discussed in previous molecular dynamics (MD) studies^17^. Furthermore, V1094 was observed to be proximal to the BH residue K830 in the closed TL state^4^. An interaction between K830 and V1094 side-chains could be counter-intuitive and possibly undervalued. However, neutralization of lysine’s positive charge through ionic interactions (such as D836) can promote hydrophobicity of the lysine side chain^55^, supporting the observed K830-V1094 interactions in the TL closed state (Supplementary Fig. 3c). Most variants in V1094 are LOF (Fig. 4c, left), consistent with disruption of K830-V1094 interaction and concomitant destabilization of the closed, active TL conformation.

Models for NTP substrate-induced TL closing remain largely untested^4,14-17^. A recent Pol II structure^51^ exhibiting an open TL state led to explicit implication of a hydrophobic pocket formed by TL residues (A1076, M1079, T1080, G1097 and L1101) and other TL proximal residues (I837, L841, V1352, V1355 and I1356) in substrate-induced TL-folding (Supplementary Fig. 3d). Q1078 recognition of the 2’-OH of a matched NTP substrate was proposed to promote release of the adjacent residue M1079 from the hydrophobic pocket, triggering TL closing^51,56^. Consistent with disruption of this observed pocket and concomitant destabilization of the inactive open TL state, A1076T, a pocket variant previously isolated as genetically GOF, conferred increased transcription activity *in vitro* (Fig. 5b). Notably, GOF phenotypes were observed for a large number of variants in pocket residues. Among them, we observed almost universal GOF phenotypes for G1097 variants, but not the extreme fitness defects found for the previously observed GOF variant G1097D. We individually phenotyped ten G1097 variants from the traditional screening and confirmed this observation (Supplementary Fig. 3e). Together, these results are consistent with the hydrophobic pocket stabilizing the inactive, open TL and providing a plausible mechanism for substrate-induced TL closing where a single residue, M1079, can act as linchpin for the entire TL through a network of interactions.

**Figure 5:**
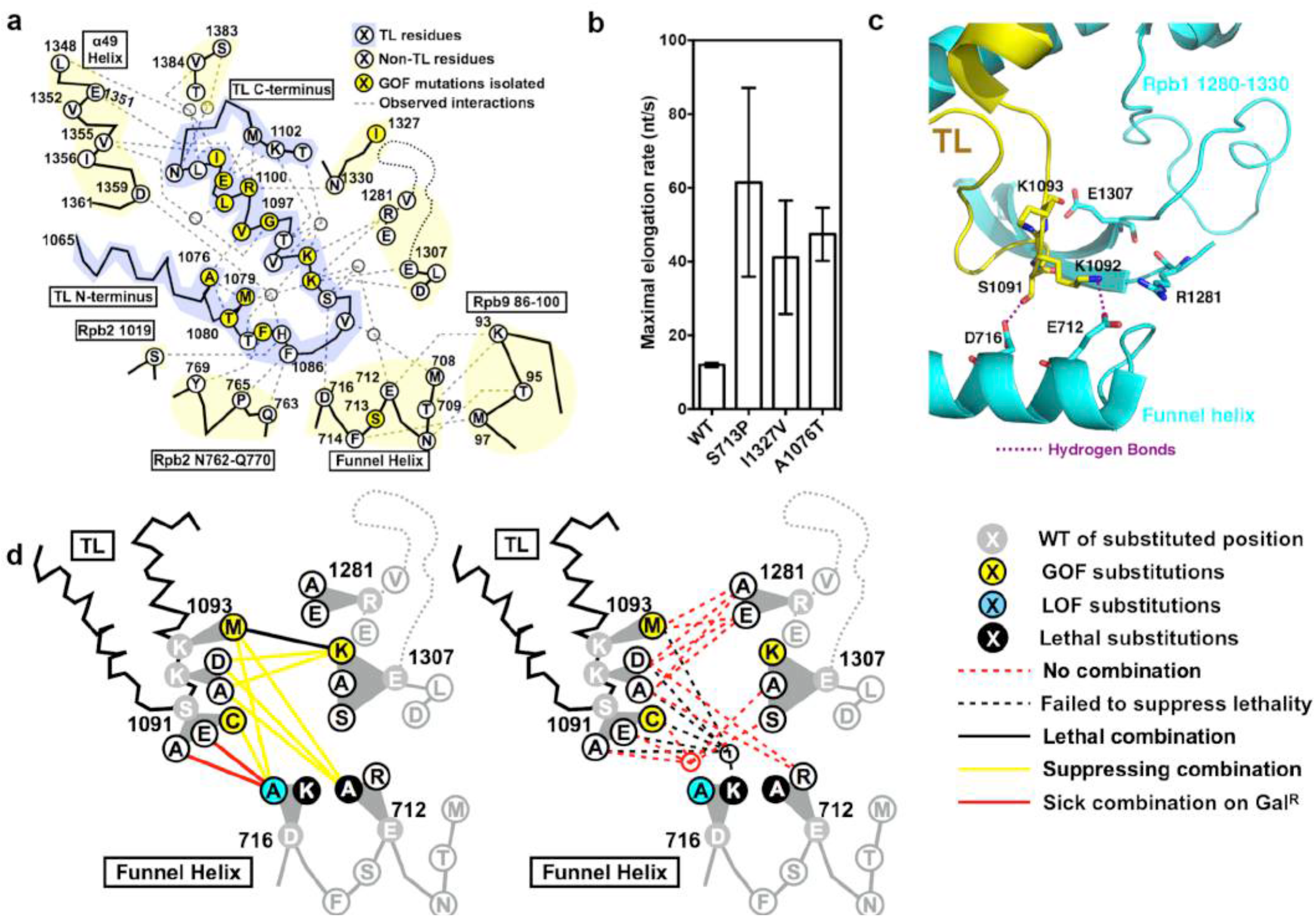
Functional contribution of TL tip and funnel helix to proper TL dynamics. Figure 5a: Observed and predicted interactions between TL and TL-proximal domains. TL schematic is shown with residues identified by single-letter amino acid code and positions of interest with positions of isolated GOF mutants color coded in yellow. Observed TL interactions with other Rpb1 domains from structures or simulation studies are shown in the grey dashed lines. Figure 5b: Maximal elongation rates of Pol II WT and genetically GOF mutants S713P, I1327V and A1076T. Figure 5c: Observed interactions between open TL tip and TL adjacent charged residues (PDB: 5C4X). Figure 5d: Genetic interactions between TL tip and proximal Rpb1 domains. Schematic of TL and the adjacent domains are shown in lines, with positions of interests shown in single-letter code. Substituted residues are shown in grey, with substituting amino acids shown in white, blue or yellow filled circles based on single mutant phenotypes (Supplementary Figure 3). Double substitution phenotypes are shown as colored lines connecting the two relevant single substitutions.

### Identification of stress conditions that alter transcription *in vivo*

GOF and LOF TL variant classes have distinct stress response profiles. In general, compared to LOF variants, GOF mutants are more sensitive to Mn^2+^, caffeine and cycloheximide yet more resistant to hydroxyurea and formamide (Fig. 4d). The allele-specific Mn^2+^ response amplified our previous observation that GOF E1103G was highly sensitive to Mn^2+^ while the LOF H1085Y was resistant to Mn^2+^, and Mn^2+^ effects on both mutants were suppressed by Mg^2+^ supplementation^52^. The TL phenotypic landscape showed that this Mn^2+^ response was general and class-specific for GOF and LOF mutants (Fig. 4d). To validate this observation, we individually analyzed seven additional variants (two LOF and five GOF) for Mn^2+^ sensitivity in the presence or absence of Mg^2+^ supplementation. Notably, all tested LOF mutants conferred Mn^2+^ resistance while all tested GOF mutants conferred Mn^2+^ hypersensitivity (Fig. 4e). Allele-specific Mn^2+^responses could be suppressed by Mg^2+^ supplementation (Fig. 4e). Mn^2+^ has been shown to stimulate transcriptional activity while compromising fidelity *in vitro*^53,54^. Our observation suggested the possibility that Mn^2+^may suppress LOF mutants by stimulating transcriptional activity yet exacerbate GOF mutants by further decreasing their already compromised transcriptional fidelity *in vivo*^11,12^. Increased Pol II catalytic activity correlates strongly with upstream transcription start site (TSS) shift *in vivo*^36,40^; therefore we assayed for TSS alterations upon Mn^2+^ treatment. Primer extension analysis at *ADH1* revealed that Mn^2+^ treatment shifted the TSS distribution upstream, and further exacerbated the upstream shift in E1103G (Fig. 4f). Deletion of *PMR1*, the golgi Mn^2+^ export channel, causes accumulation cytosolic Mn^2+ 57,58^, and can be used to alter Mn^2+^ levels apart from supplementing medium. Our prior high throughput genetic interaction analyses of Pol II mutants showed that *pmr1Δ* strongly interacts with Pol II mutants in a highly allele-specific fashion, suggesting an intimate relationship between increased cellular Mn^2+^levels and altered transcription activity. Here we find that *pmr1Δ* also shifted *ADH1* TSSs upstream (Fig. 4f). While Mn^2+^ may have other indirect affects on Pol II mutants, these observations support direct effects of Mn^2+^ on Pol II transcription activity *in vivo*, raising the possibility that other allele-specific stress conditions ( *e.g.* formamide) may also directly alter transcription *in vivo*.

### Functional contributions of the TL tip region

The TL tip region (Rpb1 1090-1096) is a random-coil region that forms an α-helical structure when the TL is closed, and helical formation has been proposed to assist TL closing^8,17,42^. Mejia *et al* characterized two *Eco* RNAP TL tip mutants I1134V and G1136S (Equivalent to *Sce* Pol II V1094 and S1096) with slightly decreased or increased transcription activity, respectively42. These results were interpreted as I1134V and G1136S substitutions decreasing or increasing the helical propensity and thus disfavoring or favoring TL closing^42^. *Sce* Pol II contains each of these variants as the WT residue, therefore individual substitutions to the *E. coli* variants (V1094I and S1096G) would be predicted to confer opposite phenotypes under the helical propensity model. However, V1094I and S1096G did not confer clear phenotypes consistent with either GOF or LOF (Fig. 4b), failing to support the helical propensity model. We asked if the proposed correlation from *Eco* RNAP studies was a general property for TL substitutions in this region, if extended to more than two substitutions. Our data, calculated from 122 variants, fail to support a general correlation between helical propensity and predicted catalytic activity for Pol II substitutions in this region (Supplementary Fig. 6a). As discussed above, V1094 may be involved in interaction with BH residue K830, and LOF in most V1094 variants may result from disrupted BH/TL coordination. Therefore, we repeated the analyses excluding V1094 variants, yet still failed to observe a correlation (Supplementary Fig. 6a). We cannot rule out contributions of helical propensity in this region to TL function; however, we did not find compelling or widespread evidence for it.

A number of recent studies have suggested potential functions of the TL tip region in regulating TL dynamics^17,51,59^. In a simulated TL closing process, positively charged K1092 and K1093 were predicted to interact with several TL-proximal residues, and some of the predicted interactions were validated by subsequent Pol II crystal structures with alternative open TL states (Fig. 5a). These interactions were proposed to stabilize the open, inactive TL state, and thus alanine (K1092A, K1093A) or charge reversing substitutions (K1092D/E, K1093D/E) were predicted to disrupt the inactive TL open state and result in GOF^17^. Contrary to this prediction, none of the above substitutions conferred GOF (Fig. 4b). Networks of residue-residue interactions near the TL tip were observed^17,51^, some of which may be functionally overlapping or redundant, adding complexity to simple models. Our previous point mutant epistatic miniarray profile (p-EMAP) studies predicted two TL-proximal mutants (S713P and I1327V) to be GOF, which we confirm here (Fig. 5b), suggesting that perturbation near the TL may interfere with native interactions, or create new ones, to destabilize the open TL. The tested variants here also extend the correlation between genetically predicted GOF and increased activity *in vitro* (Fig. 5b). Additionally, several TL tip variants with bulky side chains (K1092W, K1093Y, K1093M) conferred GOF phenotypes (Fig. 4b). Given the complexity and observation of both GOF/LOF phenotypes, we wished to further assess the functions of these residue-residue interactions.

Functional interactions among residues can be explored by the similarity between single substitution variants and the phenotypes of double mutants. We first sought evidence that variants in the potential interaction partners could confer similar GOF or LOF phenotypes. In the simulation, K1092 switched interaction partners between two funnel helix residues D716 and E712^17^, and other charged residues were either observed or simulated to interact with S1091, K1092 or K1093 (Fig. 5a). Therefore, we constructed a panel of mutants in the residues D716, E712, R1281, E1307, and D1309 for phenotypic analyses. Notably, we observed GOF phenotypes (Mn^S^ and MPA^S^) in E1307K but not E1307A, suggesting that E1307K gained an interfering interaction to destabilize the open TL state. Furthermore, we observed the Gal^R^ phenotype in D716A (Fig. 5d, Supplementary Fig. 6f), consistent with LOF. D716K and E712A were lethal (Fig. 5d, Supplementary 6b, 6c), but their defects were further explored by double mutant analyses (discussed below). Together, both GOF and LOF variants were observed in the TL tip proximal residues, consistent with roles in regulating TL dynamics.

To further dissect functional relationships, we phenotyped double mutants from potential interaction partners, and observed a number of genetic interactions (Fig. 5d, Supplementary Figure 6g-k). First, GOF and LOF mutants were mutually suppressive when combined, and most TL mutants from same biochemical class (GOF/GOF or LOF/LOF) showed additive effects (synthetically sick or lethal). The observed class-specific genetic interactions are similar to the previously reported intra-TL genetic interactions^36^, consistent with alteration of TL function in TL tip proximal variants. Furthermore, K1092A/D single substitutions did not confer transcription phenotypes, but were able to suppress the E1307K GOF phenotypes. This observed epistasis suggested that loss of K1092 relieved a putative gain of interaction in E1307K (discussed above). Finally, E712A lethality was fully suppressed by K1092A, K1092D or K1093M, adding an additional instance of epistasis. A model to explain this complex genetic relationship is that loss of native E712-K1092 interaction re-directed K1092 towards an alternative interaction or strengthened an existing interaction with D716, causing lethality. Alteration of TL tip interaction potential through K1092/1093 substitutions relieves this allele-specific effect. Taken together, the observed allele-specific and epistatic interactions between TL tip and proximal residues suggest a highly complex genetic network of residues controlling TL dynamics, and illustrate how individual residues might constrain or allow diversification of the TL through evolution.

### Functional interplay of TL and Bridge helix (BH)

The BH is a strikingly conserved structural domain of multisubunit RNA polymerases spanning the wide central cleft between polymerase “jaws”, adjacent to the active site and proximal to the TL^1,60,61^. Although BH is a straight helix in most published structures^1-6^, some *Thermus thermophilus* RNAP structures reveal a bent BH conformation proposed to support translocation^60^. This BH bending mechanism was supported by a number of simulation studies but has never been directly tested^1,24,60-62^. In the archaea *Methanocaldococcus jannaschii (Mja)* RNAP, proline substitutions at two hinge-proximal residues M808 and S824 (equivalent to *Sce* Rpb1 M818 and T834) resulted in GOF, suggesting kinking by proline substitution results in increased translocation or catalysis^62,63^. Furthermore, *Mja* GOF TL and BH mutants were not additive when combined, suggesting mutual dependence on BH and TL functions^62^.

To explore the functional consequence of BH kinking in *Sce* Pol II, we constructed and phenotyped BH mutants analogous to the characterized GOF and LOF variants in *Mja* RNAP. Notably, *Sce* T834 and other BH C-terminal hinge substitutions conferred *in vivo* phenotypes consistent with the altered transcriptional activities in *Mja* RNAP (Fig. 6a), and we directly confirmed the altered activity of T834 variants *in vitro* (Fig. 6b). In contrast, substitutions in M818, a predicted BH N-terminal hinge, showed defects deviating from expected conservation of function. M818P caused lethality, and could not be suppressed by all tested TL variants, precluding us from classifying it (Supplementary Fig. 7a). Furthermore, M818S and M818Y, although viable, did not confer any clear phenotypes (Fig. 6a). Therefore, we further assessed the functional interplay between BH and TL by double mutant analyses, including BH variants (M818S/Y, T834A/P) and TL substitutions covering a range of altered transcriptional activities (Fig. 6f-i). Notably, the GOF BH variant T834P, along with M818S and M818Y, were mutually suppressive with biochemically strong LOF TL variants (Fig. 6c, 6d, 6f), revealing both additive behavior between BH and TL for some combinations, and cryptic phenotypes for M818S/Y. The LOF BH variant T834A also suppressed GOF TL variants (Fig. 6e). However, the additive interactions (exacerbation, synthetic lethality) we observed for GOF BH and TL double mutants were in contrast to the epistasis for *Mja* RNAP^62^.

**Figure 6:**
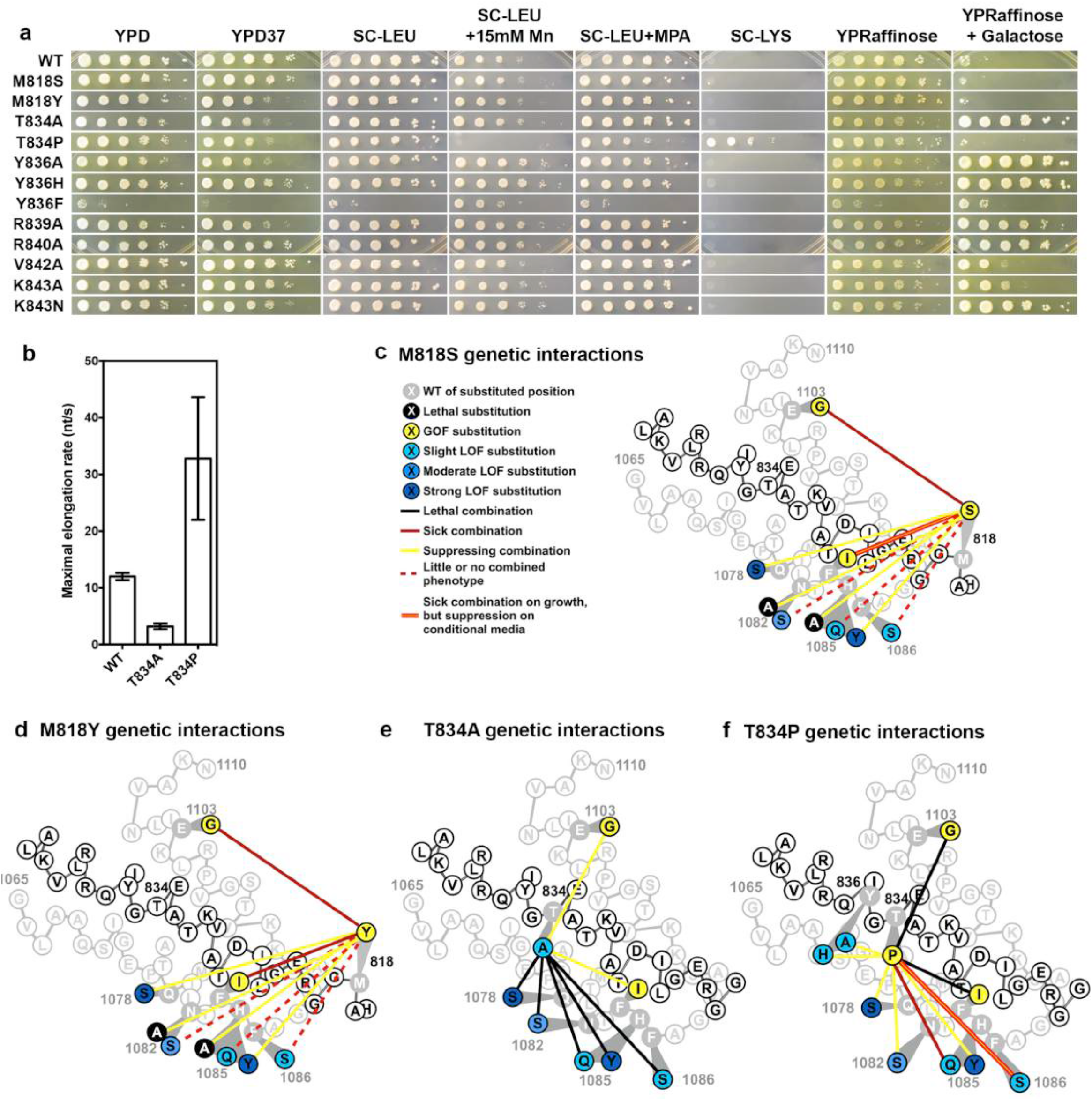
Functional interplay between TL and Bridge Helix (BH) Figure 6a: Transcription-linked phenotypes of BH single-substituted mutants. M818S and M818Y are substitutions in a predicted BH N-terminal hinge; others are substitutions in predicted BH C-terminal hinge positions or additional C-terminal substitutions. Figure 6b: Maximal elongation rate of BH variants T834A and T834P *in vitro*. Figure 6c: Genetic interactions between BH M818S and TL substitutions. M818S suppressed (yellow lines) the strong LOF TL variants (dark blue) but not the slight and moderate LOF TL variants (light blue), and M818S was synthetic sicker (red line) with the GOF TL variants (yellow). Figure 6d: Genetic interactions between BH M818Y and TL substitutions. Similar to M818S genetic interaction with TL variants (Figure 6c), M818Y suppressed (yellow lines) the strong LOF TL variants (dark blue) but not the slight and moderate LOF TL variants (light blue), and M818Y was synthetic sick (red line) with GOF TL variants (yellow). Figure 6e: Genetic interactions between BH T834A and TL substitutions. T834A suppressed (yellow lines) the GOF TL variants and was synthetic lethal with all the tested LOF TL variants (blue). Figure 6f: Genetic interactions between BH T834P and TL or BH. Similar to M818 variants (Figure 6c, 6d), T834P suppressed the strong and moderate LOF TL variants (dark blue) but was synthetic sick with the slight LOF TL variants (light blue), while synthetic lethal with the GOF TL variants (yellow). T834P was also suppressed (yellow) by two LOF BH mutants Y836A/H.

Multiple lines of evidence suggested additional, specific defects exist in BH mutants, beyond simple cooperation with the TL. First, M818P lethality could not be suppressed by all tested TL variants (Supplementary Fig. 7a), which cover a wide range of transcriptional activities. Second, suppression between BH and TL mutants of different biochemical classes (GOF/LOF) was partial and not as strong as previously observed intra-TL suppression. Third, GOF M818S, M818Y and T834P variants appeared to exhibit activity-dependent genetic interactions with TL variants. BH GOF variants suppressed strong LOF TL variants Q1078S and H1085Y but failed to suppress, or even exacerbated slightly LOF TL variants H1085Q and F1086S (Fig. 6c, 6d, 6f), consistent with conditional epistasis, where GOF activity of BH variants can suppress either specific TL variants or otherwise exert their effects in specific contexts. Finally, recent modeling studies predicted BH residue Y836 to assist Pol II forward translocation^64^ by interacting with the DNA:RNA hybrid. Y836A/H conferred Gal^R^ phenotypes, consistent with LOF and compromised translocation (Fig. 6a). Notably, GOF T834P was suppressed by Y836A/H (Fig. 6f), consistent with T834P conferring TL independent fast translocation defect, suppressible by Y836A/H.

### Context dependence of TL function

We previously observed that E1103G, a GOF allele in *Sce* Pol II, caused LOF in Pol I, highlighting divergent contributions of active site residues in different enzymatic contexts^41^. We also observed that Pol I TL and L1081M (in this study) were functionally impaired in the Pol II context. We next sought to determine the functional compatibility of other evolutionary TL variants in the *Sce* Pol II context, using our fitness and phenotypic landscape (Fig. 7). Most tested evolutionary TL variants did not confer fitness defects, though several did (Fig. 7a). Furthermore, some variants, although compatible for general growth, conferred transcription-related phenotypes and could be further classified by phenotypic landscape (Fig. 7b). These observations further demonstrate that the evolution of TL function is shaped by likely epistasis between the TL and proximal domains.

**Figure 7:**
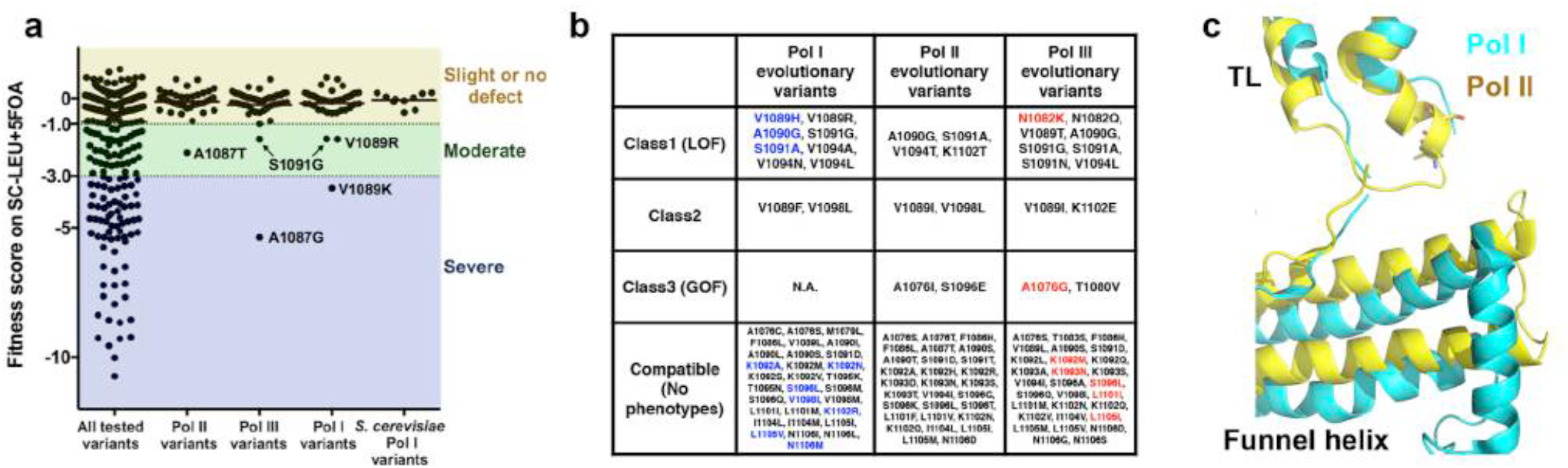
Phenotypic analyses of evolutionary variants reveal context dependent function of many TL residues. Figure 7a: General growth fitness defects of the TL single-substituted variants observed in TL across Pol I, II, III evolution including 38 Pol II, 42 Pol I and 42 Pol III amino acid variants relative to *Sce* Pol II. Figure 7b: Evolutionary TL variants in three mutant classes from the TL phenotypic landscape (Figure 4a, 4b). Existing variants from *Sce* Pol I are colored in blue, and existing variants from *Sce* Pol III are colored in red.*Sce* Pol I has three substitutions (V1089H, A1090G and S1091A) that cause LOF in the Pol II context;*Sce* Pol III has one substitution (A1076G) classified as GOF and one substitution (N1082K) classified as LOF. Figure 7c: Difference in the funnel helix positioning (relative to TL) in Pol I and Pol II. Cartoon representation of TL/funnel helix from Pol I and Pol II are shown in cyan and yellow, respectively (PDB: 5C4J and 2VUM).

We next asked what substitutions might underlie the large difference in compatibility of the *Sce* Pol I TL (versus the *Sce* Pol III TL) within Pol II^41^. From our phenotypic landscape, although many individual *Sce* Pol I and Pol III TL substitutions appeared to be compatible, functionally impairing variants were identified (Fig. 7b). The yeast Pol III TL contains Pol II GOF (A1076G) and LOF (N1082K) variants, both of which hypothetically could be mutually suppressive, resulting in close to WT activity in the Pol II context^41^. The Pol I TL contains three Pol II LOF mutations (V1089H, A1090G and S1091A). The net incompatibility of Pol I TL is consistent with additive defects of the three LOF variations, given that most TL LOF combinations show additive effects^36^. Since three evolutionarily observed variants with LOF phenotypes were all localized in the TL tip, we examined the difference between Pol I and Pol II structures for the TL tip proximal domains^51,65^. The Pol I funnel helix appears to impose less constraint than the Pol II funnel helix (Fig. 7c), suggesting that Pol I controls its TL with a distinct network of interactions. In all, our mutational data, together with the recent Pol I crystal structure, reveals enzyme-specific mechanisms to control a highly conserved domain at the heart of eukaryotic transcription.

## Discussion

The ability of the TL to fold into multiple conformations and the dynamic conversion between these states are critical for its functions. Previous studies from us and others demonstrate that TL function is delicately balanced, such that perturbations result in either increased or decreased catalytic activity and altered translocation dynamics. Distinct consequences for transcriptional activity manifest *in vivo* as what we term LOF and GOF phenotypes. In this study, we have advanced our genetic framework with which to dissect Pol II mechanisms. From our phenotypic landscape, we assessed the functional contributions of almost all TL residues to fitness in *S. cerevisiae* under multiple conditions. Our data indicate that both intra-TL interactions and TL interactions with nearby domains ( *e.g.* funnel helix) are critical for TL function. This conclusion is also supported by recent work on Rpb9 organizing the TL indirectly through an Rpb1 TL-adjacent loop^59^, interactions between the TL and F-loop regions in bacteria^30^, and predictions of TL proximal variants as GOF from our previous pEMAP analysis^40^ (validated in this study). Our system allows efficient analysis of a large number of variants to evaluate accumulating computational^17^ and structural^51^ predictions for interactions within the TL and from without.

The major function of the TL is to link substrate recognition to catalysis, and is also proposed to gate translocation such that translocation probability is linked to phosphodiester bond formation. Critical to this recognition is that a substrate be positioned correctly by base-pairing to the DNA template, and that the 2’-OH allows NTPs to be selected over 2’-dNTPs by the TL residue Q1078^4,9,27^. We have proposed that the Q1078-substrate interaction releases the adjacent M1079 from its intra-TL hydrophobic pocket, and to trigger the TL closing^51^. In this study, we show a great number of variants from the pocket residues A1076, M1079, G1097, L1101 causing GOF, providing evidence that disruption of the hydrophobic pocket destabilizes the open, inactive TL state, resulting in GOF. Additionally, while the TL shows incredibly high evolutionary conservation for a number of residues, prior work indicated alteration of ultra-conserved residues in different RNA polymerases could have distinct effects, suggesting the importance of the evolved context within each enzyme^11,36,41^. Here, we evaluate many evolutionarily observed eukaryotic TL variants in the *Sce* Pol II system, and discover a number of functionally impaired TL variants. Our results highlight that TL proximal domains may impose constraint and possibly also allow functional diversification in the molecular evolution of highly conserved TL by epistatic interactions.

The characterization of the unexpectedly healthy H1085L variant clouds the issue of how H1085 functions in substrate selection and catalysis. H1085 interacts with the substrate NTP through salt bridge and hydrogen bond^4^, and previous simulations with limiting H1085 variants predicted the hydrogen bonding to be critical for maintaining substrate interaction^66^. The discovery of H1085L argues that productive substrate interactions may be supported by entirely different chemistry, although we cannot rule out the possibility that H1085L redirects substrate interactions to an alternative residue. Furthermore, H1085 variants may have multiple defects in NAC, such as substrate selection^11^, catalysis^11,52^, intrinsic cleavage^52^ and PPi release^19,20^, and whether or not H1085 or analogous residues act as a general acid remains controversial in different RNAPs ^4,7,9,52,67^. Function of H1085L in all of these steps remains to be determined, but its phenotype suggests that function of H1085 as a general acid may be entirely bypassed.

The established TL phenotypic landscape can be further explored to study intra- and inter TL epistasis. First, whether individual TL residues work collaboratively or independently to ensure balanced TL dynamics and proper function is an open question. Some TL residues can be functionally overlapping acting at similar steps, or functionally discrete acting at distinct steps. For example, combination of LOF mutations in Q1078, N1082 and a TL-proximal residue N479 resulted in non-additive genetic interaction, suggesting functionally overlapping roles of these residues. In contrast, combination of variants from Q1078 (or N1082) and H1085 resulted in exacerbation or synthetic lethality, suggesting independent functions^36^. Coupled with structures of partially folded TL states, these genetic studies support the functional distinction between NIR residues and a multi-step TL folding model for the promotion of catalysis^36^. Here, we have identified many more predicted GOF and LOF TL variants (Fig. 4b), some of which are predicted to confer epistatic interactions ( *e.g.* F1086 and V1089). We expect the phenotypic landscape of a multiply-substituted TL library to be extremely informative for understanding functional relationship between TL residues.

Second, the TL phenotypic landscape is an extremely sensitive readout for assessing active site re-arrangement. Transcription is under control by many factors, some of which may alter the Pol II active site conformations, though few studies directly address these possibilities. Initiation factors and Pol II TL mutants confer similar alterations in transcription start site selection, consistent with initiation factors functioning through the Pol II active site and altering the efficiency of Pol II catalysis during initiation^40,52,68^. Furthermore, TL may communicate with other Pol II sites, such as the RNA exit channel or clamp domain^35^, or in direct competition with external factors, such as TFIIS^32^. Perturbations of this communication may alter TL dynamics and cause allele-specific genetic interactions (Fig. 5, Fig. 6). Specifically, an external perturbation by a relevant factor or Pol II TL distant domain may show epistasis or synergy only with specific TL alleles of a class (either LOF or GOF), whereas a non-interacting factor may not. Finally, similar perturbation of the TL phenotypic landscape by different factors would suggest functional similarity between them, thus clustering of phenotypic landscape changes upon different perturbations is expected to provide valuable insight.

The TL phenotypic landscape, along with our previous work^36^, illustrates a strategy of utilizing *in vivo* genetic reporters or stress response profiles to distinguish a large number of mutants with distinct *in vivo* defects. As discussed above, the phenotypic landscape sheds light on functional contribution of TL residues to its dynamics, to the mechanism of catalysis and to the evolutionary constraints of the TL sequence and function. The phenotypic landscape strategy expands the current scope of existing deep mutational scanning studies^43-46^, and can be generalized to study most, if not all, yeast proteins.

## Acknowledgments

We gratefully acknowledge Stefan Heiderich for technical assistance during the Slonomics library synthesis. We thank members of Kaplan lab, Steve Lockless, Qiuyan Shao, and Kun Ping Lu for helpful discussion and feedback on the manuscript. This work was supported by National Institutes of Health grant R01GM097260 and Welch Foundation Grant A-1763 to CDK, and by National Institutes of Health Grant R01GM036659 to Roger Kornberg during the initial stages of this project. We also acknowledge NSF REU site grant DBI-0851611 for supporting O.E.

## Online Methods

### Yeast strains, Media and Plasmids

All yeast strains are derived from a *GAL2+* derivative of S288C strain^69^. Genotypes and phenotypes of the yeast strains are in Supplementary file 1. Standard yeast media and the media for assessing established transcription phenotypes are as described previously^36^. For studies with 15mM Caffeine (Sigma), 150mM Hydroxyurea (Sigma), 5mM and 15mM MnCl2 (Sigma), 0.5M NaCl (EMD), 3% formamide (JT Baker), 6% ethanol (KOPTEC), 0.07μg/mL cycloheximide (Sigma), 10mM HCl (EMD), 10mM NaOH (EMD), 10μg/mL Benomyl (Sigma), each compound was added to the minimal SC-Leucine medium at the indicated concentration from concentrated stock solutions.

Detailed description of plasmids is in Supplementary file 1, and complete sequences of plasmids are available upon request. For studies involving individual analyses of Pol II mutants, site-directed mutagenesis was performed via the Quikchange strategy from Stratagene. All mutagenized regions have been verified by sequencing before sub-cloning into pRS315-derived plasmids, as previously described^36^.

### Genetic and biochemical analyses of individual Pol II mutants

Phenotypic analyses of individual Pol II mutants were performed by plasmid shuffling assays, with viable mutants further subjected to standard plate phenotyping. Each mutant in a pRS315-derived plasmid (*CEN LEU2*) was transformed into CKY283 (*rpb1Δ::CLONATMX*, pRP112 *RPB1 CEN URA3*). Transformants (Leu^+^) were patched on SC-LEU plates and subsequently replica plated to SC-LEU+5FOA (1mg/mL) to assay complementation ability upon loss of the *RPB1 CEN URA3* plasmid. Experimental details are as previously described^11,36^. Saturated cultures from single colonies of viable and shuffled Pol II mutants were subject to 10-fold serial dilution and spotting on indicated phenotyping media, as described in various previous reports^11,36^.

Pol II enzymes were purified via a tandem-affinity tag (TAP) protocol derived from^70^ with modifications described in^11^. Transcription elongation reactions were performed with Pol II elongation complexes assembled on a nucleic acid scaffold, in a procedure described in^11^ with slight modifications in the amount of Pol II and nucleic acids as described in^51^. For each enzyme, elongation assays were performed with 25μM, 125μM, 500μM and 750μM NTPs (each of ATP, GTP, CTP, UTP), and maximal elongation rates were extracted exactly as previously described^11^.

*ADH1* transcription start site selection was analyzed by primer extension. In brief, indicated strains were grown in YPD until mid-log phase (~1*10^7^ cells/mL), and diluted with YPD with 10mM MnCl_2_ or equal volume of H_2_O. Total RNA was extracted as described^71^, and 30μg of total RNA was subject to primer extension analysis, following a protocol derived from^72^ with modifications described in^36^.

### High-throughput phenotypic analyses of the TL variants library

The TL variant library was synthesized by Sloning Biotechnology (now MorphoSys) with well-characterized TL variants excluded (specified in Fig. 1B) using a building block approach^47,48^. The TL variant library was transformed into CKY283 via a gap-repair strategy as previously described^40^. In brief, the amplified TL variant library with flanking sequence was transformed into CKY283 together with a linearized pRS315-derived plasmid (*CEN LEU2*) containing *rpb1* deleted for the TL (*TLΔ*) and linearized at the deletion junction, allowing *in vivo* homologous recombination and resulting in a library of complete *rpb1* genes containing TL variants. The gap-repaired TL variants (Leu^+^) were titered and plated at 200-300 colonies per plate to reduce inter-colony growth competition, and Leu^+^ colonies were subsequently replica-plated to SC-LEU+5FOA (1mg/mL), and subsequently to additional selective and control media. Three independent biological replicate screens were performed. In each replicate, we pooled 6000 to 12000 colonies. Each cell pool was subjected to genomic DNA extraction and TL amplification by emulsion PCR. Amplification of the TL region was performed using Micellula DNA Emulsion & Purification (ePCR) Kit (Chimerx) per manufacturer’s instructions. To minimize amplification bias, each sample was amplified in a 15-cycle ePCR reaction, purified and subject to another 13-15 cycle scale-up ePCR reactions. The two-step ePCR amplification protocol ensured sufficient yield of DNA for NGS sequencing while minimizing perturbation of the allele distribution in the DNA pool. The amplified samples were subject to Illumina HiSeq 2500 sequencing, and on average over 2 million reads were obtained from each replicate of a sample, with high reproducibility and minimal perturbation of the mutant distribution within the TL variant library (Supplementary Fig. 1d).

Allele frequency was subsequently measured by deep sequencing of the TL amplicons. To identify the mutations that were present for each set of paired-end reads, a codon-based alignment algorithm was developed to align each paired-end read set in which the overlapping substrings from both flanking regions agreed perfectly to the WT sequence. The purpose of our approach was to identify real variants using an expected set of mutant codons used in the programmed library synthesis from sequencing errors. A dynamic programming algorithm was applied so that an exact match of three letters was assigned a positive score, a mismatch of at least one letter in a codon was assigned a negative score, and the insertion or deletion of either one, two or three letters was assigned a constant negative score. The allele frequency was subsequently calculated from the mapped reads, and the phenotypic score of each TL variant was calculated by allele frequency change (normalized to WT) under each condition, as below:

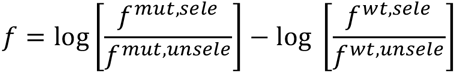

Mutants with less than 200 reads in the transformed pool (SC-LEU) and allele frequency changes assessed from less than 50 reads from both conditions were excluded from further analyses. Median values from three independent biological replicates were used for fitness and phenotype scoring. Fitness score cutoff for lethality was estimated based on fitness scores (on SC-LEU and 5FOA) of 163 known viable TL and 16 known lethal mutants. Hierarchical clustering for generating phenotypic landscape was performed by Gene Cluster 3.0 using centered correlation^73^. Figures displaying structural information were generated using Pymol (https://www.pymol.org/).

### Evolutionary analyses

Eukaryotic RNA polymerase large subunit sequences were obtained from BLAST using *Sce* Rpb1 (Pol II), *Sce* Rpa190 (Pol I), and *Sce* Rpo31 (Pol III) sequences as queries. Sequences were assigned to Pol I, II, or III based on highest similarity when compared to each of the three query sequences, with prokaryotic sequences further filtered out. Multiple sequence alignments (MSAs) were generated by first applying CD-HIT^74^ to cluster sequences so that the identity between sequences in different clusters was less than 90%, then applying MUSCLE^75^ to obtain an alignment that contains one representative sequence from each cluster. The TL conservation score was generated using Jalview^76^ and plotted as a heatmap using Gene-E (http://www.broadinstitute.org/cancer/software/GENE-E/index.html).

